# A cooperative network of molecular “hot spots” highlights the complexity of LH3 collagen glycosyltransferase activities

**DOI:** 10.1101/841486

**Authors:** Antonella Chiapparino, Francesca De Giorgi, Luigi Scietti, Silvia Faravelli, Tony Roscioli, Federico Forneris

## Abstract

Hydroxylysine glycosylations are collagen-specific post-translational modifications essential for maturation and homeostasis of fibrillar as well as non-fibrillar collagen molecules. Lysyl hydroxylase 3 (LH3) is the only human enzyme capable of performing two chemically-distinct collagen glycosyltransferase reactions using the same catalytic site: inverting beta-1,O-galactosylation of hydroxylysines and retaining alpha-1,2-glycosylation of galactosyl hydroxylysines.

Here, we used structure-based mutagenesis to show that both glycosyltransferase activities are strongly dependent on a broad cooperative network of amino acid side chains which includes the first-shell environment of the glycosyltransferase catalytic site and shares features with both retaining and inverting enzymes. We identified critical “hot spots” leading to selective loss of inverting activity without affecting the retaining reaction. Finally, we present molecular structures of LH3 in complex with UDP-sugar analogs which provide the first structural templates for LH3 glycosyltransferase inhibitor development.

Collectively, our data provide a comprehensive overview of the complex network of shapes, charges and interactions that enable LH3 glycosyltransferase activities, expanding the molecular framework for the manipulation of glycosyltransferase functions in biomedical and biotechnological applications.

## 1. INTRODUCTION

Collagens are the most abundant proteins in the human body and are highly conserved from sponges to mammals (Luther et al, 2011; Myllyharju & Kivirikko, 2004). The different oligomeric architectures and roles of collagen molecules strongly depend on a variety of post-translational modifications, including proline and lysine hydroxylations, as well highly specific glycosylations of hydroxylated lysines (HyK) (Cummings, 2009; Myllyharju & Kivirikko, 2004). The disaccharide present on HyK contains a highly conserved glucosyl(α−1,2)-galactosyl(β−1,O) glycan moiety, whose identity was discovered in the late sixties (Spiro & Spiro, 1971; Spiro, 1967; Spiro, 1969). Monosaccaridic galactosyl-(β−1,O) HyK have been identified as well, as a result of catabolic reactions carried out by the collagen α-glucosidase, an enzyme highly specific for the disaccharide present on collagenous domains. The role for this enzyme is to localize collagen in the glomerular basement membrane (Hamazaki & Hamazaki, 2016; Sternberg et al, 1982; Sternberg & Spiro, 1980). The spread of glycosylation largely depends on collagen type (Bornstein & Sage, 1980; Spiro, 1969; Terajima et al, 2014), on the functional area inside tissues (Moro et al, 2000; Schofield et al, 1971; Toole et al, 1972), on the developmental stage (Rautavuoma et al, 2004; Sipila et al, 2007) and on disease states (Lehmann et al, 1995; Salo et al, 2008; Tenni et al, 1993). Although extensively studied, the precise mechanisms of collagen glycosylation and their biological relevance in collagen homeostasis have remained poorly understood.

The identity and exquisite stereochemistry of the Glc(α−1,2)-Gal(β−1,O)-HyK-linked carbohydrate supports the idea that at least two distinct enzyme types are needed to fully incorporate these complex post-translational modifications on collagen molecules (Hennet, 2019). The first reaction indeed requires an inverting-type galactosyltransferase (GalT) acting on HyK, whereas the subsequent glucosylation is catalyzed by a retaining-type glucosylgalactosyltransferase (GlcT). Multifunctional lysyl hydroxylase 3 (LH3) was the only enzyme identified as possessing lysyl hydroxylase activity as well as GalT and GlcT activities *in vitro* (Wang et al, 2002a). The glycosyltransferase activities are specific of LH3, as highly homologous LH1 and LH2a/b are not capable of catalyzing these reactions (Heikkinen et al, 2000). Conversely, *in vivo* studies have demonstrated that decreased LH3 protein levels and/or pathogenic mutations in LH3 GT domain, exclusively impair the GlcT activity (Ewans et al, 2019; Salo et al, 2008; Savolainen et al, 1981). This occurs secondarily to the LH3 p.Asn223Ser, which introduces an additional glycosylation site within the enzyme’s GT domain leading to an osteogenesis imperfecta-like phenotype (Salo et al, 2008); and in the recently identified LH3 p.Pro270Leu, which results in a Stickler-like syndrome with vascular complications and variable features typical of Ehlers-Danlos syndrome and Epidermolysis Bullosa (Ewans et al, 2019). Mouse studies have also shown that only the LH3 GlcT activity is indispensable for the biosynthesis of collagen IV and formation of the basement membrane during embryonic development (Rautavuoma et al, 2004; Ruotsalainen et al, 2006), consistent with the presence of additional collagen galactosyltransferases. Two genes encoding for O-galactosyltransferases (*GLT25D1* and *GLT25D2*) were recently identified (Perrin-Tricaud et al, 2011; Schegg et al, 2009). It is of interest that GLT25D1 and LH3 were proposed to act in concert on collagen molecules (Schegg et al, 2009; Sricholpech et al, 2011; Yamauchi & Sricholpech, 2012). Studies on osteosarcoma cell lines which produce large amounts of fibrillar collagens, showed that the simultaneous deletion of *GLT25D1* and *GLT25D2* resulted in growth arrest due to lack of glycosylation, further indicating that the GalT activity of LH3 might not be as essential as its GlcT activity (Baumann & Hennet, 2016).

These data support the existence of a specific and highly conserved machinery for collagen O-glycosylation, and sustain the hypothesis that *in vivo* the entire collagen glycosylation machinery may involve distinct proteins and protein complexes for GalT and GlcT reactions. This raises the intriguing question of how this highly conserved process is spatiotemporally regulated at the molecular level. Our current understanding of collagen glycosyltransferases is however restricted to three-dimensional structures of human LH3 in complex with UDP-sugar donor substrates (Scietti et al, 2018) and to few mutagenesis studies focusing on the main hallmarks of Mn^2+^-dependent glycosyltransferase catalysis (Wang et al, 2002a; Wang et al, 2002b).

Here, we combine site-directed mutagenesis scanning with biochemistry and structural biology to characterize the glycosyltransferase activities of human LH3. Our data highlight an overall distribution of “hot spots” around the extended glycosyltransferase cavity of LH3, critically involved in both GalT and GlcT functions, and very few amino acid residues capable of selectively abolishing transfer of galactose to HyK without affecting GlcT activity. Finally, we also identify and characterize UDP-sugar substrate analogs acting as inhibitors of LH3 glycosyltransferase activities.

Together, our results provide insights into the LH3 glycosyltransferase activities and expand the available structural framework for the development of collagen GalT/GlcT inhibitors. These insights will assist with the manipulation of LH3 protein functions and donor substrate specificity in biomedical applications.

## RESULTS

### Features and roles of the non-conserved LH3 “glycoloop”

The LH3 N-terminal GT domain shares its fold with Mn^2+^-dependent GT-A glycosyltransferases, encompassing a UDP-donor substrate binding cavity stretched towards a GalT/GlcT catalytic pocket (Scietti et al, 2018). We firstly inspected the structural organization of the UDP binding cavity to identify signatures for LH3 glycosyltransferase activity by comparing the amino acid sequences for the residues surrounding the UDP-sugar donor substrate with those of GalT/GlcT-inactive LH1 and LH2a/b isoforms. We found that nearly all residues involved in Mn^2+^ and UDP-sugar binding are conserved in orthologs, with the exception of Val80, becoming Lys68 in LH1 and Gly80 in LH2a/b (Fig 1). The presence of a different amino acid side chain surrounding the donor substrate cavity led us to consider whether this could be a discriminating functional feature among the GT domains in LH enzymes. In LH3, Val80 is located in the middle of a flexible “glycoloop” (Gly72-Gly87), not visible in the electron density of the ligand-free LH3 structure and stabilized upon UDP-substrate binding (Scietti et al, 2018). Within this glycoloop, residue Val80 is in close proximity to the ribose ring of the UDP-sugar donor substrates. We hypothesized that introduction of a large, positively charged residue such as Lys in LH1, or alterations due to complete removal of side chain steric hindrance such as Gly in LH2 could lead to inability of binding donor substrates. We therefore generated the LH3 Val80Lys and Val80Gly mutants. Both enzyme variants were found to be folded based on analytical gel filtration and differential scanning fluorimetry (DSF), and showed lysine hydroxylation activity comparable to wild-type LH3 (supplementary Fig S1). Conversely, the mutation resulted in significant reduction of both GalT and GlcT activities when the reaction was carried out in presence of both donor and acceptor (gelatin) substrates (Fig 2, supplementary Table S1). Considering that wild-type LH3 is also capable of activating donor UDP-sugar substrates and release UDP in absence of the acceptor collagen substrate (“uncoupled” activity, as defined in (Scietti et al, 2018)), we investigated the impact of the Val80Lys and the Val80Gly mutations also in absence of acceptor substrates. In this case, the experiments yielded minor, but significant reduction in the enzymatic activities between wild-type and mutant LH3 (Fig 2, supplementary Table S1), indicating that the Val80 residue might be involved in the productive positioning of the donor substrate during transfer of the glycan moiety to the acceptor substrate, rather than stabilizing the UDP moiety in the catalytic pocket.

**Figure 1:**
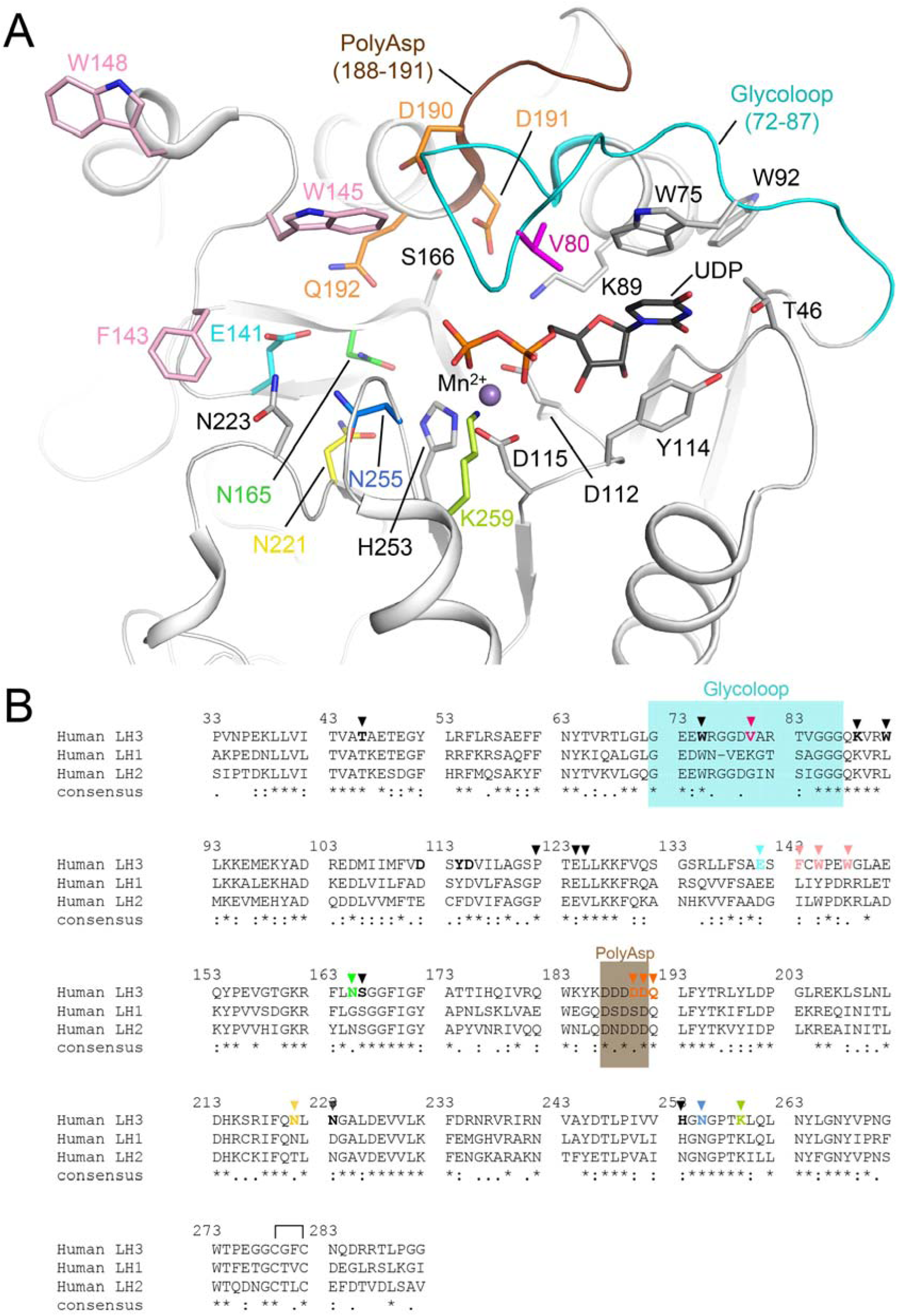
Features of the LH3 glycosyltransferase (GT) domain. (A) Cartoon representation of the LH3 GT domain (PDB ID: 6FXR) showing the key residues shaping the catalytic site as sticks. The PolyAsp motif (brown) and the Glycoloop (cyan) involved in binding of UDP-sugar donor substrates are shown. The residues implicated in the catalytic activity and investigated in this works are colored, while the residues depicted in grey have already been shown to be essential in Mn^2+^ (purple sphere) and UDP (black sticks) coordination. (B) Sequence alignment of human LH1, LH2 and LH3 GT domains, highlighting similarities and differences in the amino acid residues within the active site. Residues shown in Fig 1A are indicated with a triangle. Colored boxes indicate the PolyAsp motif (brown) and the Glycoloop (cyan).

**Figure 2:**
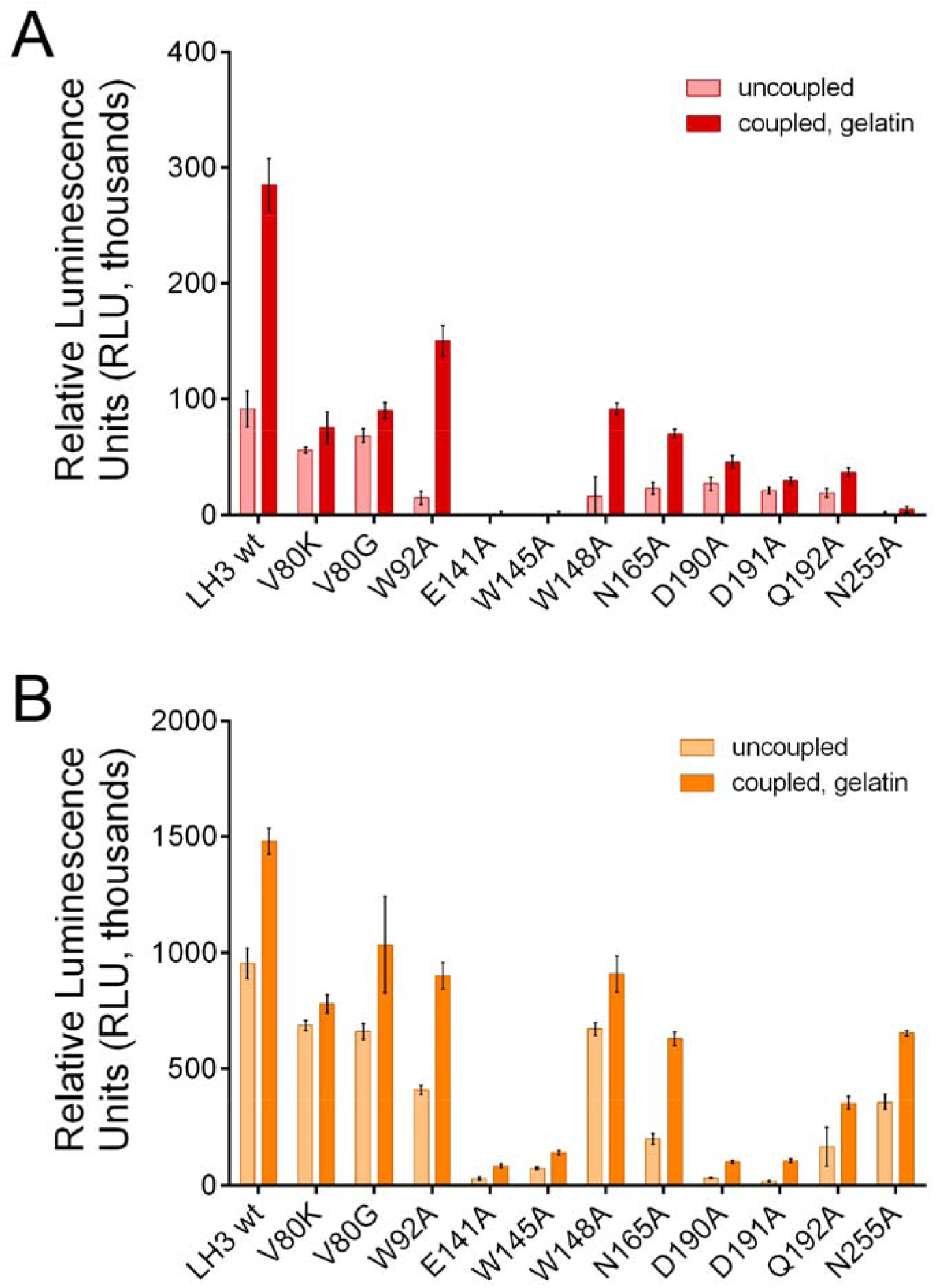
Evaluation of the effect of LH3 GT domain mutations in the GT site on glycosyltransferase activities. Evaluation of the GalT activity (A) and GlcT activity (B) of LH3 mutants compared to the wild-type. Each graph shows the enzymatic activity detected in absence (i.e., “uncoupled”) or in presence of gelatin, used as acceptor substrate. The plotted data are baseline-corrected, where the baseline was the background control. Error bars represent standard deviations from average of triplicate independent experiments.

To further rationalize the implications of LH3 Val80 in GalT and GlcT activities, we crystallized and solved the 3.0-Å resolution structure of the Val80Lys mutant in complex with Mn^2+^, and also obtained its 2.3-Å resolution structure in presence of both Mn^2+^ and the UDP-Glc donor substrate (supplementary Table S2). Overall, both structures superimpose almost perfectly with wild-type LH3 for all domains (supplementary Fig S2). The structure of the LH3 Val80Lys mutant bound to Mn^2+^ appeared identical to that of wild-type LH3. In both structures, the glycoloop containing Val/Lys80 could not be modelled in the electron density due to its high flexibility (supplementary Fig S2B). On the other hand, the side chain of the Val80Lys residue could be modelled unambiguously in the experimental electron density of the UDP-donor substrate bound structure (Fig 3A). Despite the increased steric hindrance, the mutated Lys80 residue adopted a conformation compatible with the simultaneous presence of the UDP-Glc in the catalytic cavity. However, similar to what observed for wild-type LH3, the glycan moiety of UDP-Glc was completely flexible and therefore not visible in the electron density (Fig 3A). Collectively, these data are consistent with the alteration introduced by the Val80Lys mutation, impacting partially on the LH3 glycosyltransferase catalytic activities.

**Figure 3.**
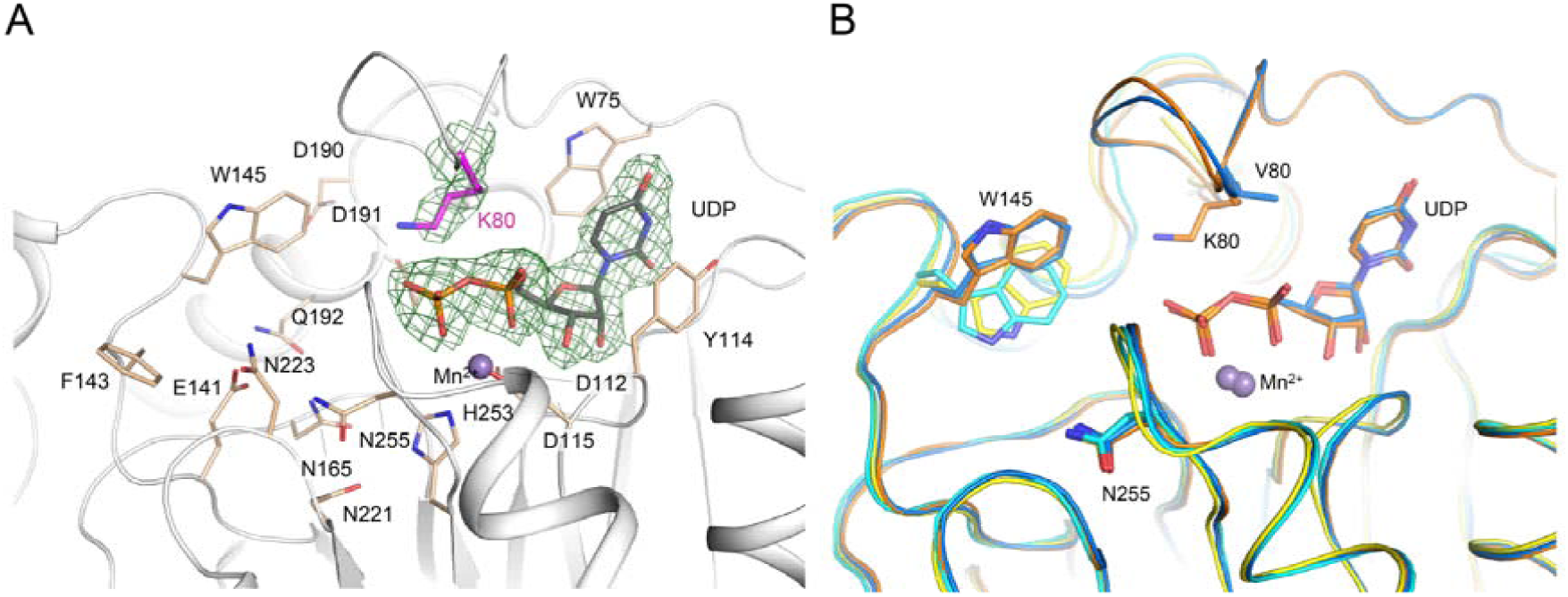
Structural characterization of the LH3 Val80Lys mutant. (A) Crystal structure of the LH3 Val80Lys mutant in complex with UDP-glucose and Mn^2+^. Electron density is visible for the mutated lysine and the UDP donor substrate (green mesh, 2*F*_*o*_-*F*_*c*_ omit electron density map, contoured at 1.3 σ). Catalytic residues shaping the enzyme cavity are shown as sticks, Mn^2+^ is shown as purple sphere. Consistent with what observed in the crystal structure of wild-type LH3, the glucose moiety of the donor substrate is not visible in the experimental electron density. (B) Superposition of wild-type and Val80Lys LH3 crystal structures in substrate-free (cyan and yellow, respectively) with UDP-glucose bound (marine and orange, respectively) states. Notably, the conformations adopted by the side chain of Trp145 upon ligand binding are consistent in the wild-type and in the mutant enzyme. As the glycoloop is flexible in substrate-free structures, the side chains of Val/Lys80 are visible only in the in UDP-sugar bound structures.

The glycoloop is a structural feature found exclusively in the GT domains of LH enzymes. It incorporates Trp75, a residue whose aromatic side chain stabilizes the uridine moiety of the donor substrate and, together with residue Tyr114 of the DxxD motif (a distinguishing feature of LH3 GT domain, shared among LH enzymes, (Scietti et al, 2018)) “sandwiches” the donor substrate in an aromatic stacking environment (Fig 1A). Both residues are critical for the LH3 GalT and GlcT enzymatic activities (Scietti et al, 2018). The conformation adopted by the LH3 glycoloop in the presence of UDP-donor substrates is however not accompanied by other significant structural changes in surrounding amino acids, with the exception of minor rearrangements of distant residue Trp92 (not conserved in other LH isoenzymes (Fig 1)), whose bulky side chain rearranges pointing towards the aromatic ring of Trp75. Prompted by this observation, we mutated this residue to alanine and found that the presence of this variant did not alter the folding of the enzyme (supplementary Fig S1A-B) nor its LH enzymatic activity (supplementary Fig S1C). Conversely, the mutant showed 40% decrease for both GalT and GlcT activities in presence of donor and acceptor substrate compared to LH3 wild-type (Fig 2, supplementary Table S1); the impact of the Trp92Ala mutation on reactions in absence of acceptor substrate seemed to affect both activities at similar levels, with lower residual GalT (30%) compared to GlcT (40%) (Fig 2, supplementary Table S1). These findings suggest that LH3-specific long-range interactions in the GT domain may contribute to the productive conformations of the glycoloop in donor substrate-bound states.

In UDP-sugar bound structures, the glycoloop contacts a poly-Asp sequence (Asp188-Asp191, Fig 1A); this sequence is partially conserved in LH isoforms lacking glycosyltransferase activities (Fig 1B). Mutations of Asp190 and Asp191 were reported to affect the glycosyltransferase enzymatic activities of LH3 (Wang et al, 2002b). Based on LH3 crystal structures, such behaviour is expected, since residues Asp190 and Asp191 point towards the GalT/GlcT catalytic cavity. Interestingly, superposition of LH3 molecular structures with GT-A fold glycosyltransferases show that Asp191 caps the N-terminal end of an α-helix in a highly conserved position, with functional relevance in both retaining (Flint et al, 2005; Persson et al, 2001) and inverting enzymes (Charnock & Davies, 1999; Pedersen et al, 2000) (supplementary Fig S3, Table 1). We designed and generated individual alanine mutants for both LH3 Asp190 and Asp191, and found that both variants were compatible with folded and functional LH enzymes (supplementary Fig S1). When tested for GalT and GlcT activity, both these mutants caused severe impairment in activation of donor UDP-sugar (as shown by the strong reduction in uncoupled activity to less than 5% using UDP-Glc as substrate) as well as in transfer of the sugar moiety to the acceptor substrate (Fig 2, supplementary Table S1). However, none of these mutations resulted in a complete inactive LH3 GalT nor GlcT glycosyltransferase. Collectively, these data point towards an extended involvement of the residues of the poly-Asp repeat, and in particular Asp190 and Asp191, in both the positioning and recognition of donor or acceptor substrates.

**Table 1:**
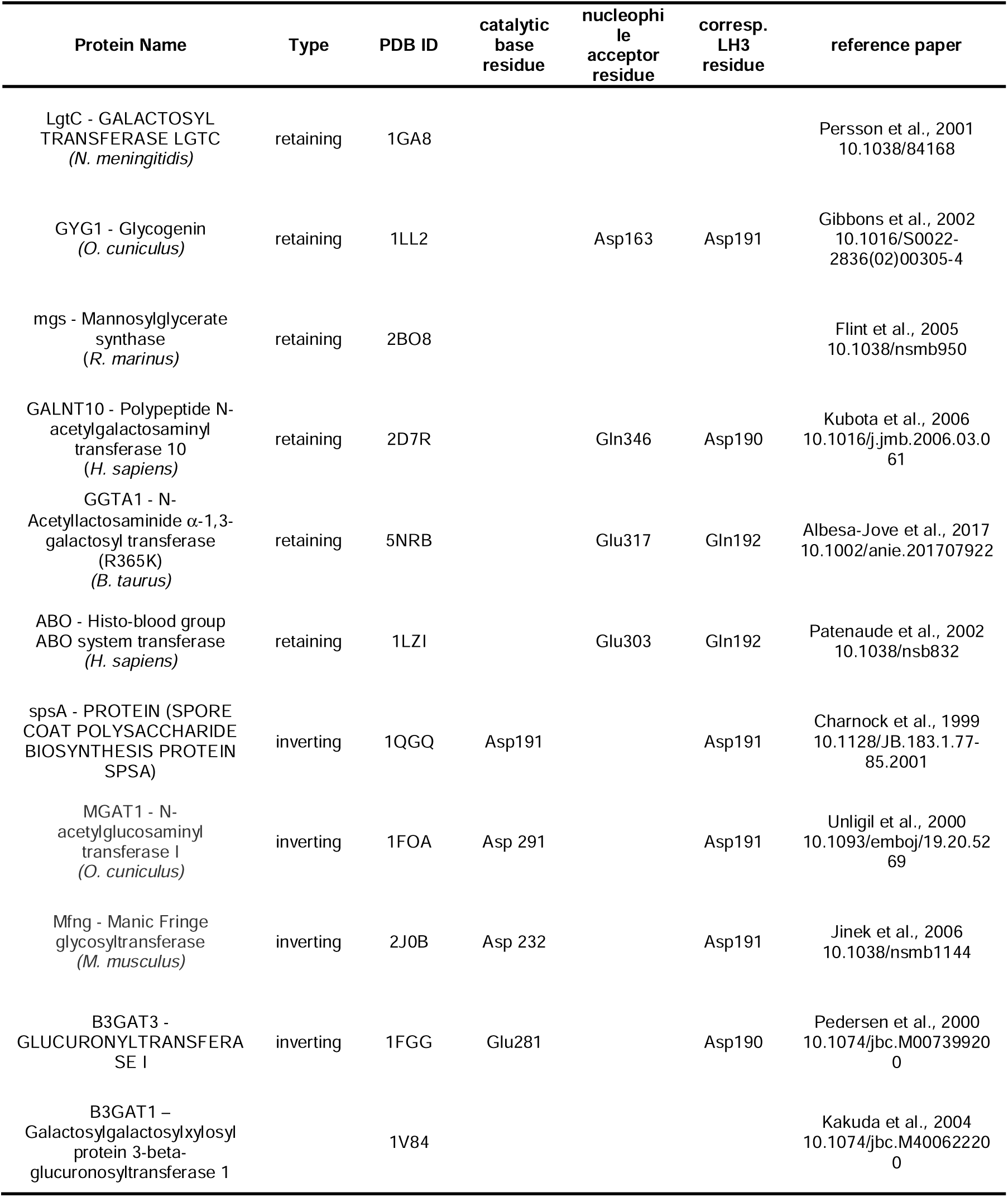
List of glycosyltransferase enzymes used for comparisons with human LH3. The list includes the indication of the catalytic bases and nucleophile residues as proposed in the original papers describing the various glycosyltransferases, with the corresponding residue number in human LH3 based on structural superpositions.

### LH3 shares features with both retaining and inverting glycosyltransferases

After investigating the amino acid residues involved in stabilization of the UDP moiety of donor substrates, we focused on another group of residues within the GT catalytic pocket, opposite to the putative position of the flexible sugar rings of the same substrates (Fig 1A). Many LH3 residues shaping this part of the glycosyltransferase catalytic cavity matched catalytic amino acids found in other GT-A type glycosyltransferases (Fig 1A, supplementary Fig S3) (Ardevol et al, 2016; Lairson et al, 2008). In particular, LH3 Trp145, a residue located in one of the loops of the GT domain uniquely found in LH3, was previously suggested as a possible candidate for modulation of LH3 GalT and GlcT activities. This residue was found to adopt different side chain conformations in substrate-free and substrate-bound structures, affecting the shape and steric hindrance of the enzyme’s catalytic cavity (Scietti et al, 2018); interestingly, nearly identical conformational changes were observed when comparing substrate-free and substrate bound LH3 Val80Lys structures (Fig 3B). Mutating the LH3 Trp145 residue into alanine strongly reduced both GalT (6% residual) and GlcT (10% residual) enzymatic activities (Fig 2, supplementary Table S1), without affecting other enzyme’s properties (supplementary Fig S1). This supports previous hypotheses of a “gating” role for Trp145 in the GT catalytic cavity, assisting the productive positioning of sugar moieties of donor substrates for effective transfer during catalysis. Comparison with molecular structures of other glycosyltransferases (including distant homologs) highlighted that most structurally-related enzymes manage to position aromatic side chains from different structural elements of their fold in their catalytic cavities. Such structural arrangement is reminiscent to that of Trp145 in LH3, but relies on completely different structural features of the glycosyltransferase domain. In particular, similar aromatic residues were found in other glycosyltransferases such as Tyr186 in LgtC from *Neisseria meningitidis*, Trp314 in the N-acetyllactosaminide α-1,3-galactosyl transferase GGTA1, Trp300 in the histo-blood group ABO system transferase, and Trp243 and Phe245 in the two glucoronyltransferases B3GAT3 and B3GAT1, respectively (Table 1, supplementary Fig S3). This supports glycosyltransferases being highly versatile enzymes, displaying an impressive structural plasticity to carry out reactions characterized by a very similar mechanism on a large variety of specific donor and acceptor substrates.

Our previous structural comparisons of ligand-free and substrate-bound LH3 highlighted the additional possibility of a concerted mechanism involving conformational changes of a non-conserved aromatic residue located on the LH3 surface (Trp148, Fig 1), together with Trp145 (Scietti et al, 2018). To investigate such possibility, we mutated Trp148 into alanine. The mutant enzyme was also found to behave like wild-type LH3 in folding and LH activity (supplementary Fig S1). Glycosyltransferase assays showed that this variant had reduced GalT (30% residual) and GlcT (50% residual) activities compared to the wild-type, in presence of both donor and acceptor substrates (Fig 2, supplementary Table S1). Despite less pronounced alterations compared to those observed when mutating Trp145, these data support possible synergistic mechanisms between long-range acceptor substrate recognition on the enzyme’s surface and conformational rearrangements in the enzyme’s catalytic site.

Molecular structures of LH3 in complex with UDP-Glc and Mn^2+^, showed weak electron density near the UDP pyrophosphate group partially compatible with glycan donor substrates, likely representative of multiple conformations simultaneously trapped in the substrate binding cavity (Scietti et al, 2018). We explored the LH3 catalytic cavity in its proximity, looking for additional amino acids potentially critical for catalysis. In particular, we searched for residues carrying carboxylic or amide side chains, thus capable of acting as candidate catalytic nucleophiles for the formation of a (covalent) glycosyl-enzyme intermediate prior to glycosylation of the acceptor substrate (Gloster, 2014).

In retaining-type glycosyltransferases belonging to the GT-6 family, a conserved glutamate has been found to act as a nucleophile (supplementary Fig S3) (Albesa-Jove et al, 2017; Coutinho et al, 2003; Gomez et al, 2012; Lombard et al, 2014; Patenaude et al, 2002). In LH3 structures, we noticed that residues Gln192, Asn165 and Glu141 point towards the cavity that accommodates the glycan moiety (Fig 1A). We generated Ala mutations of all these residues, obtaining in all cases folded functional LH enzymes (supplementary Fig S1). When probed for GalT and GlcT activity, we found that both the Asn165Ala and the Gln192Ala mutants were less efficient, but still capable, in activating UDP-donor substrates and performing sugar transfer to acceptor substrates. Conversely, the Glu141Ala mutant was completely deprived of both GalT and GlcT activities (Fig 2, supplementary Table S1). These data suggest Glu141 as essential for catalysis and the surrounding negatively charged pocket composed of Asp190, Asp191, Gln192, Asn165 comprising a broad network of amino acids which may concertedly assist the LH3 glycosyltransferase activity.

Proximate to Glu141 in the GalT/GlcT cavity, residue Asn255 is the closest amino acid to the UDP phosphate-sugar bond. Despite being fully conserved in LH isoforms (Fig 1B), this residue is not found in any GT-A-type glycosyltransferases with known structures to date. The side chain of Asn255 consistently points to a direction opposite to the donor substrate in all LH3 structures (Fig 3B, supplementary Fig S2B). We wondered whether the side chain amide group might be involved in catalysis, possibly through recognition of acceptor substrates given the conformation displayed by this side chain. Surprisingly, LH3 Asn255Ala mutants showed that their GalT activity was completely abolished, whereas the GlcT enzymatic activity was reduced to 50% (Fig 2, supplementary Table S1); the protein was properly folded and showed LH activity comparable to LH3 wild-type (supplementary Fig S1). These results identify Asn255 as a possible critical discriminating residue for the two glycosyltransferase activities of LH3, and rule out possible functions for this residue as catalytic nucleophile for retaining-type glycosyltransferase mechanisms, given its major impact restricted to the GalT catalytic activity.

### Pathogenic LH3 mutations in the LH3 GT domain affect protein folding

Recently a new pathogenic LH3 mutation, Pro270Leu, has been identified and mapped at the interface of the AC and GT domain (Ewans et al, 2019). This residue localizes in a loop which is critical for shaping the GT cavity, although given its position it is unlikely to play direct roles in catalysis. To better understand the impact of this pathogenic mutation on LH3 enzymatic activity, we generated a Pro270Leu mutant. In this case, due to very low expression levels, we could not reliably carry out any *in vitro* investigations. Considering the high reproducibility associated to recombinant production of a large variety of LH3 point mutants, this result may indicate that this mutation is likely to severely impact the overall enzyme stability rather than its enzymatic activity, resulting in extremely low protein expression levels *in vitro* and likely also *in vivo*.

### Molecular structures of LH3 in complex with UDP-sugar analogs provide insights on how glycan moieties are processed inside the LH3 catalytic cavity

A frequent limitation associated to molecular characterizations of glycosyltransferases is the high flexibility of the donor substrate glycan moiety within the catalytic cavity. Such limitation becomes even more relevant when the enzyme is capable of processing UDP-sugar molecules in absence of acceptor substrates, such as in the case of LH3. Considering our previous (Scietti et al, 2018) and current co-crystallization results, we wondered whether free UDP, the product of the enzymatic reaction, could remain bound in the LH3 GT domain with the same efficiency as physiological donor substrates even after processing. We therefore compared the binding of free UDP and donor UDP-sugars using DSF and detected a thermal shift of 3.5 °C for free UDP, compared to a 2-2.5 °C shift using UDP-sugar substrates (Fig 4A). These results suggested that free UDP may bind to LH3, likely with even higher affinity than UDP-glycan substrates, and that the GalT and GlcT reactions may therefore be affected by product inhibition. To our surprise, the increase in thermal stability did not correlate with efficient trapping of the reaction product in LH3 molecular structures. Independently from the UDP concentration used in co-crystallization and soaking experiments, we never observed any electron density for free UDP, yielding LH3 structures completely identical to ligand-free enzyme (supplementary Fig S4A).

**Figure 4.**
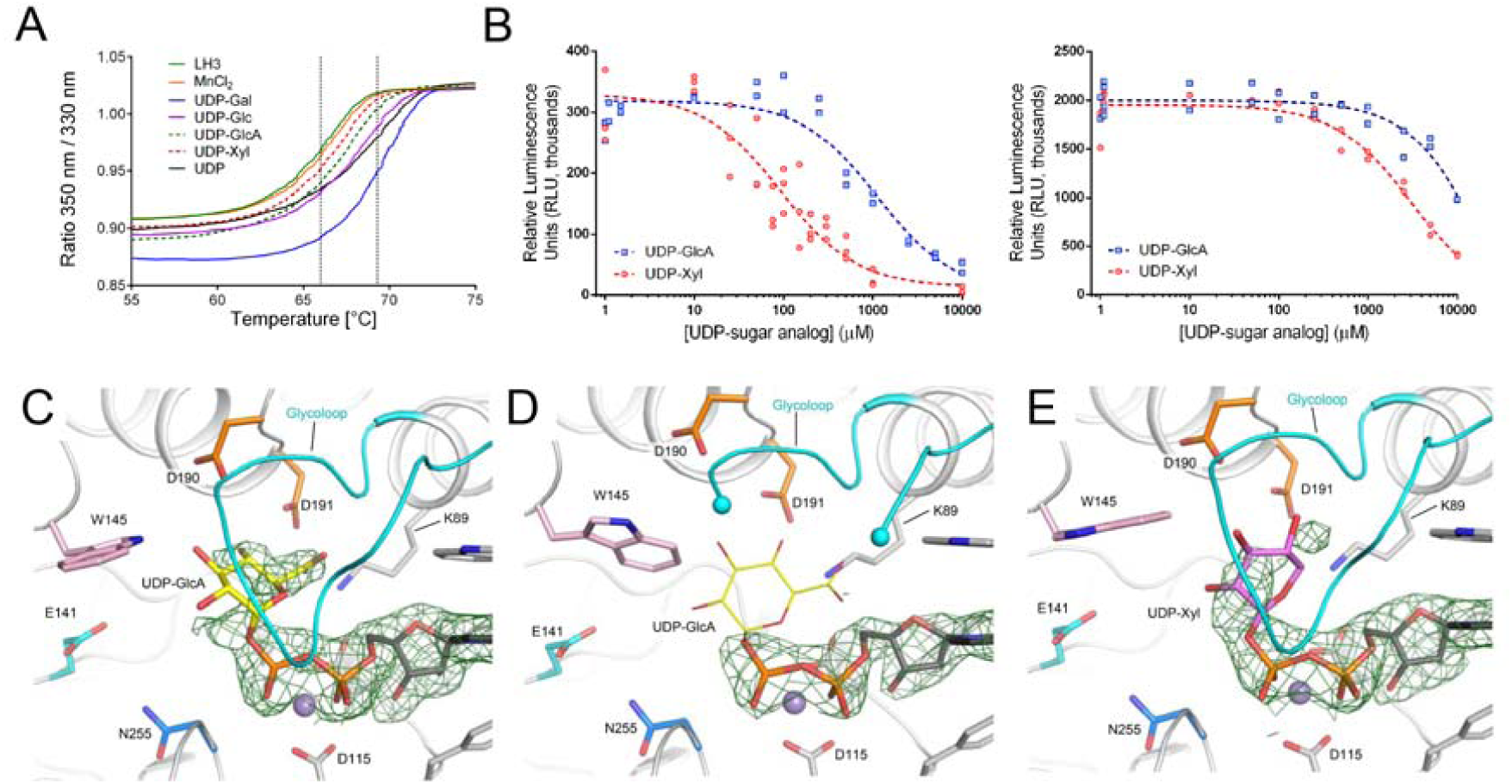
Characterization of UDP-sugar analogs. (A) Thermal stability of LH3 wild-type (solid green) using differential scanning fluorimetry (DSF) in presence of various Mn^2+^ and several UDP-sugars. A prominent stabilization effect is achieved in presence of the biological donor substrates UDP-galactose (solid blue), UDP-Glucose (solid purple) and free UDP (solid black). A milder stabilization effect is also obtained with UDP-xylose (red dash) and UDP-glucuronic acid (green dash). (B) Evaluation of GalT and GlcT enzymatic activities of LH3 in the presence of increasing concentrations of UDP-GlcA or UDP-Xyl. (C) Crystal structure of LH3 wild-type in complex with Mn^2+^ and UDP-glucuronic acid shows clear electron density for UDP (2*F*_*o*_-*F*_*c*_ omit electron density maps, green mesh, contour level 1.2 σ). The glucuronic acid (shown in yellow) can be modelled even if with partial electron density. (D) Crystal structure of the LH3 Val80Lys mutant in complex with Mn^2+^ and UDP-glucuronic acid. Whereas the UDP backbone can be modelled in the electron density (black sticks) (2*F*_*o*_-*F*_*c*_ omit electron density maps, green mesh, contour level 1.2 σ), in this case no electron density is present for the glucuronic acid (shown in yellow). In addition, the portion of the glycoloop containing the mutated lysine is flexible from residue 79 to 83 (shown as cyan spheres). (E) Crystal structure of LH3 wild-type in complex with Mn^2+^ and UDP-xylose. Similar to UDP-GlcA, UDP shows clear electron density (2*F*_*o*_-*F*_*c*_ omit electron density maps, green mesh, contour level 1.2 σ), whereas partial density is shown for the xylose moiety (shown in pink).

A report from Kivirikko and colleagues (Kivirikko & Myllyla, 1979) suggested that UDP-glucuronic acid (UDP-GlcA) could act as competitive inhibitor of collagen glycosyltransferases. Based on that, UDP-GlcA was used to isolate LH3 from chicken embryos preparation (Myllyla et al, 1977; Wang et al, 2002a). However, no follow-up biochemical studies could be found in the literature. We used DSF and luminescence-based GalT and GlcT activity assays to investigate whether and how UDP-GlcA could affect LH3 enzymatic activity. DSF showed that UDP-GlcA indeed binds weakly to LH3, resulting in a thermal shift of 1-1.5 °C (Fig 4A), highlighting limited stabilization compared to UDP-glycan substrates and free UDP. Enzymatic assays also confirmed the competitive inhibition displayed by this molecule (Fig 4B), with IC_50_ values in the millimolar range (supplementary Table S1). We also successfully co-crystallized and determined the 2.2-Å resolution crystal structure of wild-type LH3 in complex with Mn^2+^ and UDP-GlcA (supplementary Table 2), and found that the inhibitor could efficiently replace UDP-sugar donor substrates in the substrate cavity (Fig 4C). We observed additional electron density for the glucuronic acid moiety of the inhibitor in the enzyme’s catalytic cavity, however this density could not be interpreted with a single inhibitor conformation. Nevertheless, analysis of the experimental electron density for the glucuronic acid moiety unambiguously showed that the inhibitor adopts a “bent” conformation: the glycan moiety is deeply buried in the enzyme’s catalytic cavity proximate to residues Lys89, Asp190, Asp191, but distant from the residues found critical for catalysis, including Trp145, Asn255 and Glu141 (Fig 4C), thereby leaving the remaining space in the cavity for accommodating acceptor substrates.

Considering the possible conformations adopted by the glucuronic acid moiety based on analysis of the electron density and the close proximity of the glucuronic acid moiety to LH3 Val80, we wondered whether the LH3 Val80Lys mutation could interfere with inhibitor binding. We therefore co-crystallized and solved the 2.7-Å resolution structure of LH3 Val80Lys mutant in complex with UDP-GlcA (supplementary Table S2), and surprisingly observed partial displacement of the glycoloop, for which we could not observe the typical well defined electron density present in UDP-sugar-bound wild-type LH3 structures (Fig 4D). At the same time, we could not observe improvements in the quality of the electron density for the glucuronic acid moiety, resulting even poorer than what observed in wild-type LH3 (Fig 4D). This suggests that the intrinsic flexibility of the sugar-like moiety is not influenced by specific conformations of the glycoloop, but rather by lack of specific protein-ligand interactions that could provide stabilization of the sugar ring in a unique structural arrangement.

The lack of a precise conformation for the glucuronic acid moiety observed crystal structures prompted us for a further investigation of another UDP-sugar substrate analog, characterized by lack of the carboxylic moiety of UDP-GlcA: UDP-Xylose (UDP-Xyl). Similar to UDP-GlcA, UDP-Xyl resulted to be a weak inhibitor of LH3 GalT and GlcT activities (Fig 4A-B), with IC_50_ in the high micromolar range (supplementary Table S1). The 2.0-Å resolution structure of LH3 in complex with Mn^2+^ and UDP-Xyl also showed the inhibitor bound inside the enzyme’s catalytic cavity, with weak electron density associated to the sugar moiety suggesting multiple conformations of the xylose moiety attached to UDP, similar to what observed for UDP-GlcA (Fig 4E). Taken together, these results suggest that the LH3 GT binding cavity is capable of hosting a variety of UDP-sugar substrates, and that inhibition likely depends on the reduced flexibility (and therefore increased stabilization) of the ligand within the cavity. In this respect, we could expect that UDP-sugar analogs strongly interacting with side chains proximate to the glycan moieties of UDP-GlcA and UDP-Xyl, may have the potential to become powerful inhibitors of LH3 glycosyltransferase activities.

## DISCUSSION

Glycosyltransferases are highly versatile, yet very specific enzymes. If carefully inspected, they reveal a series of recurrent features that allow their comparative characterization even in presence of very low sequence/structure conservation. LH3 has been known for long time as a promiscuous enzyme able to exploit both an inverting and a retaining catalytic mechanism *in vitro* for the specific transfer of different sugars to at least two different acceptor substrates: the HyK and the GalHyK of collagens (Myllyharju & Kivirikko, 2004). Our *in vitro* investigations highlight multiple areas surrounding the glycosyltransferase catalytic site that can be considered as critical “hot spots” for the LH3 GalT and GlcT activities: the glycoloop, the poly-Asp helix, the acceptor substrate cavity, and the region proximate to the UDP-sugar donor substrate (Fig 1A).

LH3 is the only isoform of its family found capable of glycosyltransferase activities, however the strong sequence conservation among human LH isoforms cannot be used to elucidate the key determinants for this additional function. The GT domain has features (such as the DxxD motif, the poly-Asp region, the glycoloop) not found in other glycosyltransferases, yet highly conserved within the LH family (Fig 1B). Computational homology models of homologous LH1 and LH2a/b (Scietti et al, 2019) (supplementary Fig S5) support the possibility of UDP-sugar donor binding and processing. To further investigate LH enzymes’ GT domains, we focused on the only non-matching residue present within the whole amino acid sequence directly surrounding the UDP-donor substrate. This residue (Val80 in LH3, corresponding to Lys68 in LH1 and Gly80 in LH2a/b) is located in the middle of the glycoloop, in close proximity to the ribose ring of the UDP-sugar donor substrate(s). Our data were consistent with this residue being important for catalysis, as it assists the positioning of the bound donor substrate and in particular the glycan moiety. However, the results also emphasize how sequence alterations at this site are not sufficient to justify the lack of GalT/GlcT enzymatic activities in homologous LH1/2. We therefore expanded our investigation to the second-shell environment surrounding the donor substrate, and found that non-conserved LH3 Trp92 (Leu80 in LH1, Leu92 in LH2a/b) positions its aromatic side chain in a conformation that stabilizes the entire glycoloop to facilitate the enzymatic reactions. Again, mutating this residue did not lead to full loss of the glycosyltransferase activities. This is consistent with the loss of GalT/GlcT functions in LH1 and LH2a/b being associated to a broad set of subtle alterations, possibly involving residues distant from the actual enzyme’s catalytic site, likely essential for recognition of collagen acceptor substrates.

Prompted by these observations, we expanded our investigation to the LH3 catalytic cavity expected to host the glycan moieties of the UDP-sugar donor substrates and the acceptor molecules. Residues Asp190 and Asp191 of the characteristic LH3 poly-Asp helix lie in a conserved position compared to other retaining and inverting type GT-A (supplementary Fig S3, Table 1) and our data indicate that both these carboxylate moieties are critical for efficient donor substrate activation and sugar transfer. Although residues matching these positions have been proposed to act as catalytic nucleophiles in retaining type GT-A (Flint et al, 2005; Persson et al, 2001; Wang et al, 2002b), and as catalytic bases in inverting type GT-A (Charnock & Davies, 1999; Pedersen et al, 2000), their distances and the relative orientations with respect to UDP-sugar donor substrates in the LH3 GT domain (Fig 1A) do not support this hypothesis. Nevertheless, these residues play critical roles in recognizing and assisting the proper positioning of donor UDP-sugar substrates, as shown by their proximity to glycan moieties in LH3 co-crystal structures with UDP-sugar analogs (Fig 4C-E), a feature observed also in mutagenesis studies on other GT-A retaining type glycosyltransferases (Lairson et al, 2004).

On the opposite site of the UDP-binding pocket, LH3 exhibits a non-conserved loop shaping the GT catalytic cavity bearing two aromatic residues, the Trp145 and the Trp148 that seem to act in a concerted way during catalysis, as suggested by comparisons between substrate-free and substrate-bound LH3 molecular structures (Fig 1A). Trp145 is indispensable for both GalT/GlcT activities: its conformational changes seem to respond to the presence and conformational positioning of the donor substrate inside the catalytic cavity. Although located in a loop that is uniquely found in LH3, the Trp145 side chain matches a site frequently occupied by bulky aromatic residues in other GT-A glycosyltransferases that shape a portion of the GT cavity to facilitate donor substrate processing and catalysis. This further highlights the versatility of glycosyltransferases, in which many different structural features have evolved to specifically recognize distinct donor and acceptor substrates, while preserving the ability to carry out the same catalytic reaction. The implication in catalysis of the less conserved Trp148 on the surface of the GT domain is even more intriguing: this residue seems to coordinately respond to the rearrangements of its counterpart Trp145 in the catalytic site, suggesting involvement in recognition of the acceptor substrate prior to its access into the GT cavity. Alternatively, Trp148 may contribute to long-range stabilizing interactions with collagen molecules, while they dock their HyK or GalHyK residues in the acceptor substrate site during the GalT or GlcT reactions, respectively.

Catalytic nucleophiles have been clearly identified so far only in the retaining-type glycosyltransferases belonging to the GT-6 family (Coutinho et al, 2003; Lombard et al, 2014), such as the α−1,3 galactosyltransferase (GGTA1) where a conserved glutamate is found positioned on the β−face of the donor sugar (Albesa-Jove et al, 2017; Gomez et al, 2012; Patenaude et al, 2002) (supplementary Fig S3). Conversely, extensive structural comparisons and mutagenesis experiments have been performed in the O-galactosyltransferase LgtC from *Neisseria menengitidis*, focusing on matching residue, Gln189 (Lairson et al, 2004). However, the role of this residue as catalytic nucleophile was ruled out. This site is occupied by Gln192 in LH3. This residue is next to the poly-Asp helix, distant from the sites occupied by donor substrates and in an arrangement that is not compatible with a direct role in catalysis. However, our mutagenesis data indicate that removal of the Gln192 side chain has a strong impact on LH3 glycosyltransferase activity. In close proximity, we identified two other amino acid residues potentially involved in donor substrate activation or transfer of sugar moieties to the acceptor molecule. Both Asn165 and Glu141 point directly towards the glycan moiety of the donor substrate (Fig 1A). Whilst the Asn165Ala mutation only reduced the glycosyltransferase activity by a factor of two (Fig 2, supplementary Table 1), we found that Glu141 is essential for both GalT and GlcT activities, as the Glu141Ala mutation yields results in a completely inactive LH3 glycosyltransferase (Fig 2, supplementary Table 1). In LH3, Glu141 adopts a conformation corresponding to Asp130 in the O-galactosyltransferase LgtC from *Neisseria meningitidis*, Asp125 in the O-glucosyltransferase GYG1 from rabbit, and Gln247 in the O-glucosyltransferase GGTA1 from *Bos taurus* (supplementary Fig 3, Table 1). In LH3, residue Asn255 is the closest amino acid to the UDP phosphate-sugar bond, but in crystal structures its side chain consistently points to a direction opposite to the donor substrate (Fig 1A). When inspecting the molecular structures of other GT-A glycosyltransferases, we noticed that this residue is not conserved (supplementary Fig 3). On the contrary, this residue is fully conserved among human LH isoforms (Fig 1B). Strikingly, the LH3 Asn255Ala mutant showed a complete loss of GalT activity, but partially preserved the GlcT activity. Although the significance of *in vivo* LH3 GalT activity is uncertain, such activity is clearly detectable *in vitro* (Scietti et al, 2018; Wang et al, 2002a). The ability of Asn255 to selectively abolish only LH3 GalT activity highlights how LH3 can promiscuously accept and recognize very different acceptor substrates (i.e., collagen HyK *versus* GalHyK) within the same catalytic site and carry out two glycosyltransferase reactions that rely on different mechanisms (i.e., inverting GalT and retaining GlcT). Presently, LH3 is the only known glycosyltransferase capable of such promiscuity, and our data show that it shares numerous features with both retaining and inverting glycosyltransferases. Collectively, these results suggest the intriguing possibility that LH3 is not just the ancestor of the whole LH family, but may preserve in its sequence features belonging to the evolutionary precursors of both retaining and inverting glycosyltransferases.

In addition, the present work provides a set of 3D structures of LH3 in complex with UDP-sugar analogs, which work as mild inhibitors (Fig 4). Despite the high flexibility observed for the glycan moieties of the bound molecules, the new molecular structures presented provide valuable insights for structure-based drug development of inhibitors of LH3 GalT/GlcT enzymatic activities. These molecules may give the spark to innovative therapeutic strategies against pathological conditions characterized by excess collagen glycosylations, such as osteogenesis imperfecta (Raghunath et al, 1994).

Together with the mutagenesis scanning of the entire GT catalytic site, our work provides a comprehensive overview of the complex network of shapes, charges and interactions that enable LH3 GalT and GlcT activities.

## EXPERIMENTAL PROCEDURES

### 5.1 Chemicals

All chemicals were purchased from Sigma-Aldrich (Germany) unless otherwise specified.

### 5.2 Site-directed mutagenesis

The LH3 coding sequence (GenBank accession number BC011674.2 - Source Bioscience), devoid of the N-terminal signal peptide was amplified using oligonucleotides containing in-frame 5′-BamHI and 3′-NotI (supplementary Table 3) and cloned in a pCR8 vector, that was used as a template for subsequent mutagenesis experiments. All LH3 mutants were generated using the Phusion Site Directed Mutagenesis Kit (ThermoFisher Scientific). The entire plasmid was amplified using phosphorylated primers. For all mutants the forward primer introduced the mutation of interest (supplementary Table 3). The linear mutagenized plasmid was phosphorylated prior to ligation. All plasmids were checked by Sanger sequencing prior to cloning into the expression vector.

### 5.3 Recombinant protein expression and purification

Wild-type and mutant LH3 coding sequences were cloned into the pUPE.106.08 expression vector (U-protein Express BV, The Netherlands) in frame with a 6xHis-tag followed by a Tobacco Etch Virus (TEV) protease cleavage site. Suspension growing HEK293F cells (Life Technologies, UK) were transfected at a confluence of 10^6^ cells ml^-1^, using 1 μg of plasmid DNA and 3 μg of linear polyethyleneimine (PEI; Polysciences, Germany). Cells were harvested 6 days after transfection by centrifuging the medium for 15 minutes at 1000 x *g*. The clarified medium was filtered using a 0.2 mm syringe filter and the pH was adjusted to 8.0 prior to affinity purification as previously described (Scietti et al, 2018). All proteins were isolated from the medium exploiting the affinity of the 6xHis tag for the HisTrap Excel (GE Healthcare, USA) affinity column. The purified protein was further polished using a Superdex 200 10/300 GL (GE Healthcare) equilibrated in 25 mM HEPES/NaOH, 200 mM NaCl, pH 8.0, to obtain a homogenous protein sample; peak fractions containing the protein of interest were pooled and concentrated to 1 mg mL^-1^.

### 5.4 Crystallization, data collection, structure determination and refinement

Wild-type and mutant LH3 co-crystallization experiments were performed using the hanging-drop vapor-diffusion method protocols as described in (Scietti et al, 2018), by mixing 0.5 μL of enzyme concentrated at 3.5 mg mL^-1^ with 0.5 μL of reservoir solutions composed of 600 mM sodium formate, 12% poly-glutamic-acid (PGA-LM, Molecular Dimensions), 100 mM HEPES/NaOH, pH 7.8, 500 μM FeCl_2_, 500 μM MnCl_2_, supplemented with 1 mM of the appropriate UDP-sugar analogs (UDP, UDP-glucose, UDP-glucuronic acid, UDP-xylose). Crystals were cryo-protected with the mother liquor supplemented with 20% ethylene glycol, harvested using MicroMounts Loops (Mitegen), flash-cooled and stored in liquid nitrogen prior to data acquisition. X-ray diffraction data were collected at various beamlines of the European Synchrotron Radiation Facility, Grenoble, France and at the Swiss Light Source, Villigen, Switzerland. Data were indexed and integrated using *XDS* (Kabsch, 2010) and scaled using *Aimless* (Evans & Murshudov, 2013). Data collection statistics are summarized in Suppl. Table 2. The data showed strong anisotropy and therefore underwent anisotropic cut-off using STARANISO (Tickle et al, 2018) prior to structure determination and refinement. The structures were solved by molecular replacement using the structure of wild-type LH3 in complex with Fe^2+^, 2-OG and Mn^2+^ (PDB ID: 6FXM) (Scietti et al, 2018) as search model using *PHASER* (McCoy et al, 2007). The structure was refined with successive steps of manual building in *COOT* (Emsley et al, 2010) and automated refinement with *phenix.refine* (Adams et al, 2010). Model validation was performed with *MolProbity* (Chen et al, 2010). Refinement statistics for the final models are reported in supplementary Table 2.

### 5.5 Evaluation of LH3 GalT/GlcT enzymatic activity

LH, GalT and GlcT activities were tested using luminescence-based enzymatic assays with a GloMax Discovery (Promega, USA) as described in *Scietti et al.* (Scietti et al, 2018). Minor modifications have been done for GalT/GlcT competitive inhibition assays, where 1 μL of a mixture of 250 μM MnCl_2_, 500 μM UDP-galactose (GalT assay) or UDP-glucose (GlcT assay) and increasing concentrations of either UDP-GlcA or UDP-Xyl were initially added to the enzyme and gelatin substrate to start the reactions. All experiments were performed in triplicates. Control experiments were performed in the same conditions by selectively removing LH3. Data were analyzed and plotted using the GraphPad Prism 7 (Graphpad Software, USA).

### 5.6 Differential Scanning Fluorimetry (DSF)

DSF assays were performed on LH3 wild-type and mutants using a Tycho NT.6 instrument (NanoTemper Technologies GmbH, Germany). LH3 samples at a concentration of 1 mg/mL in a buffer composed of 25 mM HEPES, 500 mM NaCl, pH 8. Binding assays were performed by incubating LH3 with 50 μM MnCl_2_ and 5 mM free UDP or UDP-sugar donor substrates or their analogs. Data were analyzed and plotted using the GraphPad Prism 7 (Graphpad Software, USA).

## ACKNOWLEDGEMENTS

We thank the European Synchrotron Radiation Facility (ESRF) and the Swiss Light Source (SLS) for the provision of synchrotron radiation facilities. We thank Dr. M. Campioni for useful discussions and M. Miao for support in crystallization experiments.

## AUTHOR CONTRIBUTIONS

AC generated and purified LH3 mutants with support from SF, carried out biochemical investigations and crystallized LH3 mutants; LS purified and crystallized wild-type LH3 in complex with UDP-sugar analogs; FDG performed LH3 enzymatic activity assays; LS, FDG and FF carried out structural refinement and analyses; TR was part of the study group focusing on LH3 pathogenic variants. FF designed the study with help from AC, FDG and LS. AC and FF wrote the paper, with contributions from all authors.

