## supplementary for "A cooperative network of molecular “hot spots” highlights the complexity of LH3 collagen glycosyltransferase activities"

**Supplementary Table S1A: evaluation of GalT and GlcT enzymatic activities (in the absence and in the presence of acceptor substrates) of the LH3 variants described in this work.**

| Variant | GalT activity | | GlcT activity | |
|---|---|---|---|---|
| | **no acceptor (RLU*1000)** | **with acceptor (RLU*1000)** | **no acceptor (RLU*1000)** | **with acceptor (RLU*1000)** |
| wild-type | 110 ± 27 | 304 ± 39 | 967 ± 93 | 1491 ± 99 |
| Val80Lys | 75 ± 1 | 94 ± 23 | 699 ± 32 | 792 ± 70 |
| Val80Gly | 87 ± 9 | 109 ± 11 | 674 ± 49 | 1047 ± 362 |
| Trp92Ala | 34 ± 9 | 169 ± 23 | 43 ± 10 | 97 ± 16 |
| Glu141Ala | 13 ± 2 | 19 ± 2 | 420 ± 27 | 913 ± 97 |
| Trp145Ala | 8 ± 2 | 19 ± 3 | 86 ± 7 | 153 ± 16 |
| Trp148Ala | 35 ± 29 | 110 ± 8 | 684 ± 40 | 921 ± 134 |
| Asn165Ala | 42 ± 7 | 89 ± 5 | 213 ± 34 | 641 ± 50 |
| Asp190Ala | 46 ± 9 | 65 ± 9 | 45 ± 3 | 115 ± 12 |
| Asp191Ala | 40 ± 3 | 49 ± 4 | 31 ± 4 | 120 ± 14 |
| Gln192Ala | 38 ± 5 | 56 ± 5 | 179 ± 118 | 367 ± 46 |
| Asn255Ala | 19 ± 3 | 24 ± 2 | 372 ± 54 | 666 ± 19 |

**Supplementary Table S1B: evaluation of the inhibitory effect of UDP-GlcA and UDP-Xyl on the GalT and GlcT enzymatic activities of wild-type LH3 in presence of acceptor substrates.**

| Inhibitor | GalT ($IC_{50}$, μM) | GlcT ($IC_{50}$, μM) |
|---|---|---|
| wild-type + UDP-GlcA | 1130 ± 370 | > 10000 |
| wild-type + UDP-Xyl | 91 ± 23 | 3170 ± 211 |

**Supplementary Table S2: crystallographic statistics for data collection, structure solution and refinement.**

[a] Values in parentheses are for reflections in the highest resolution shell.

| | LH3 + $Fe^{2+}$ + $Mn^{2+}$ + UDP | LH3 + $Fe^{2+}$ + $Mn^{2+}$ + UDP-GlcA | LH3 + $Fe^{2+}$ + $Mn^{2+}$ + UDP-Xyl | LH3 Val80Lys + $Fe^{2+}$ + $Mn^{2+}$ | LH3 Val80Lys + $Fe^{2+}$ + $Mn^{2+}$ + UDP-Glc | LH3 Val80Lys + $Fe^{2+}$ + $Mn^{2+}$ + UDP-GlcA |
|---|---|---|---|---|---|---|
| **Data Collection[a]** | | | | | | |
| X-ray source | ESRF ID30A-3 | SLS X06SA | ESRF ID23-EH2 | SLS X06SA | SLS X06SA | SLS X06SA |
| Processing programs | XDS, AIMLESS, STARANISO | XDS, AIMLESS, STARANISO | XDS, AIMLESS, STARANISO | XDS, AIMLESS, STARANISO | XDS, AIMLESS, STARANISO | XDS, AIMLESS, STARANISO |
| Space group | $C222_1$ | $C222_1$ | $C222_1$ | $C222_1$ | $C222_1$ | $C222_1$ |
| Cell parameters | a = 97.0 Å; $\alpha$ = 90°<br>b = 100.0 Å; $\beta$ = 90°<br>c = 225.2 Å; $\gamma$ = 90° | a = 98.2 Å; $\alpha$ = 90°<br>b = 100.5 Å; $\beta$ = 90°<br>c = 224.7 Å; $\gamma$ = 90° | a = 97.2 Å; $\alpha$ = 90°<br>b = 100.0 Å; $\beta$ = 90°<br>c = 224.0 Å; $\gamma$ = 90° | a = 98.0 Å; $\alpha$ = 90°<br>b = 100.8 Å; $\beta$ = 90°<br>c = 225.7 Å; $\gamma$ = 90° | a = 98.1 Å; $\alpha$ = 90°<br>b = 100.4 Å; $\beta$ = 90°<br>c = 225.2 Å; $\gamma$ = 90° | a = 98.0 Å; $\alpha$ = 90°<br>b = 99.8 Å; $\beta$ = 90°<br>c = 224.5 Å; $\gamma$ = 90° |
| Wavelength (Å) | 0.968 | 1.000 | 0.873 | 1.000 | 1.000 | 1.000 |
| Resolution (Å) | 48.84-2.30 (2.38-2.30) | 49.10-2.20 (2.26-2.20) | 48.79-2.40 (2.49-2.40) | 49.14-3.00 (3.18-3.00) | 49.02-2.30 (2.38-2.30) | 49.00-2.70 (2.83-2.70) |
| Total reflections | 230588 (21551) | 430059 (32322) | 237715 (25682) | 93694 (15275) | 251674 (15330) | 197473 (22212) |
| Unique reflections | 48288 (4455) | 56604 (4537) | 43005 (4500) | 22529 (3578) | 49068 (4243) | 30477 (3911) |
| CC1/2[b] | 0.997 (0.512) | 0.999 (0.860) | 0.992 (0.586) | 0.986 (0.490) | 0.998 (0.461) | 0.996 (0.580) |
| Redundancy | 4.8 (4.8) | 7.6 (7.1) | 5.5 (5.7) | 4.2 (4.3) | 5.1 (3.6) | 6.5 (5.7) |
| Mean I/$\sigma$(I) | 8.4 (0.9) | 11.8 (0.5) | 5.5 (0.7) | 4.5 (0.7) | 6.7 (0.8) | 9.4 (1.3) |
| Completeness (%) | 98.7 (99.7) | 99.8 (98.6) | 99.9 (99.9) | 99.0 (98.5) | 98.7 (94.0) | 99.5 (98.3) |
| $R_{sym}$[b] | 0.118 (1.476) | 0.104 (2.566) | 0.209 (1.831) | 0.227 (3.045) | 0.100 (1.351) | 0.138 (1.312) |
| $R_{pim}$[c] | 0.086 (1.107) | 0.060 (1.554) | 0.147 (1.274) | 0.186 (2.479) | 0.071 (1.155) | 0.088 (0.902) |

[b] $R_{sym} = [\Sigma_{hkl}\Sigma_j \mid I_{hkl,j} - \langle I_{hkl}\rangle \mid ] / [\Sigma_{hkl}\Sigma_j I_{hkl,j}]$, where I is the observed intensity for a reflection and <I> is the average intensity obtained from multiple observations of symmetry-related reflections.

[c] $R_{pim} = [\Sigma_{hkl} (1/(n-1))^{1/2} \Sigma_j \mid I_{hkl,j} - \langle I_{hkl}\rangle \mid ] / [\Sigma_{hkl}\Sigma_j I_{hkl,j}]$ where I is the observed intensity for a reflection and <I> is the average intensity obtained from multiple observations of symmetry-related reflections.

| | LH3 + $Fe^{2+}$ + $Mn^{2+}$ + UDP | LH3 + $Fe^{2+}$ + $Mn^{2+}$ + UDP-GlcA | LH3 + $Fe^{2+}$ + $Mn^{2+}$ + UDP-Xyl | LH3 Val80Lys + $Fe^{2+}$ + $Mn^{2+}$ | LH3 Val80Lys + $Fe^{2+}$ + $Mn^{2+}$ + UDP-Glc | LH3 Val80Lys + $Fe^{2+}$ + $Mn^{2+}$ + UDP-GlcA |
|---|---|---|---|---|---|---|
| **Refinement** | | | | | | |
| $R_{work}/R_{free}{}^{c}$ | 0.1943/0.2458 | 0.1982/0.2411 | 0.1852/0.2159 | 0.2079/0.2412 | 0.1701/0.2268 | 0.1895/0.2278 |
| Number of atoms: | 5874 | 6293 | 6019 | 5732 | 5938 | 5893 |
| Protein | 5646 | 5754 | 5754 | 5665 | 5736 | 5716 |
| Ligands | 73 | 100 | 105 | 67 | 92 | 104 |
| Solvent | 155 | 439 | 160 | - | 110 | 73 |
| Average B-factor $(Å)^2$ | 35.83 | 36.51 | 31.45 | 42.33 | 44.80 | 43.73 |
| Protein | 35.73 | 35.91 | 31.16 | 42.07 | 44.54 | 42.99 |
| Ligands | 56.66 | 58.25 | 53.94 | 63.99 | 64.67 | 91.24 |
| Solvent | 29.78 | 39.40 | 26.84 | - | 41.54 | 33.94 |
| **Structure quality** | | | | | | |
| RMS bond lengths (Å) | 0.006 | 0.005 | 0.003 | 0.004 | 0.009 | 0.003 |
| RMS bond angles (°) | 0.78 | 0.84 | 0.58 | 0.70 | 1.05 | 0.71 |
| **Ramachandran stats** | | | | | | |
| Favored (%) | 97.5 | 97.1 | 97.0 | 95.9 | 96.3 | 96.5 |
| allowed (%) | 2.5 | 2.7 | 2.9 | 3.8 | 3.6 | 3.3 |
| outliers (%) | 0.0 | 0.2 | 0.1 | 0.3 | 0.1 | 0.2 |
| PDB ID | 6TE3 | 6TES | 6TEC | 6TEU | 6TEX | 6TEZ |

[c] $R_{free}$ values are calculated based on 5% randomly selected reflections, selected prior to STARANISO correction as recommended by the software developers (Tickle et al, 2018).

**Supplementary Table S3: list of oligos used for mutagenesis**

| Oligonucleotide Name | Sequence (5'→3') |
|---|---|
| Forward LH3 Val80Lys | AAGGCTCGAACAGTTGGTGGAGGAC |
| Reverse LH3 Val80Lys | ATCACCCCCTCGCCACTCCTC |
| Forward LH3 Val80Gly | GAGCTCGAACAGTTGGTGGAGGAC |
| Reverse LH3 Val80Gly | CATCACCCCCTCGCCACTCC |
| Forward LH3 Trp92Ala | GCATTAAAGAAGGAAATGGAGAAATACG |
| Reverse LH3 Trp92Ala | CCGGACCTTCTGTCCTCCACC |
| Forward LH3 Glu141Ala | CGAGCTTCTGCTGGCCCGAGTG |
| Reverse LH3 Glu141Ala | CTGCAGAGAAGAGCAGGCGGCTG |
| Forward LH3 Trp145Ala | GCACCCGAGTGGGGGCTGGC |
| Reverse LH3 Trp145Ala | GCAGAAGCTCTCTGCAGAGAAGAGC |
| Forward LH3 Trp148Ala | GCAGGGCTGGCGGAGCAGTAC |
| Reverse LH3 Trp148Ala | CTCGGGCCAGCAGAAGCTCTC |
| Forward LH3 Asn165Ala | GCTTCTGGTGGATTCATCGGTTTTGC |
| Reverse LH3 Asn165Ala | GAGGAAGCGCTTCCCCGTGC |
| Forward LH3 Asp190Ala | CTGACCAGCTGTTCTACACACGGC |
| Reverse LH3 Asp190Ala | CGTCATCATCCTTGTACTTCCACTGG |
| Forward LH3 Asp191Ala | CTCAGCTGTTCTACACACGGCTC |
| Reverse LH3 Asp191Ala | CGTCGTCATCATCCTTGTACTTCCAC |
| Forward LH3 Gln192Ala | GCGCTGTTCTACACACGGCTCTAC |
| Reverse LH3 Gln192Ala | GTCGTCGTCATCATCCTTGTACTTCC |
| Forward LH3 Asn255Ala | GCCGGTCCCACTAAGCTGCAGC |
| Reverse LH3 Asn255Ala | TCCATGGACCACAATGGGGAGCGTG |
| Forward LH3 Pro270Leu | TCAATGGCTGGACTCCTGAGGG |
| Reverse LH3 Pro270Leu | GGACGTAGTTTCCCAGGTAGTTGAG |

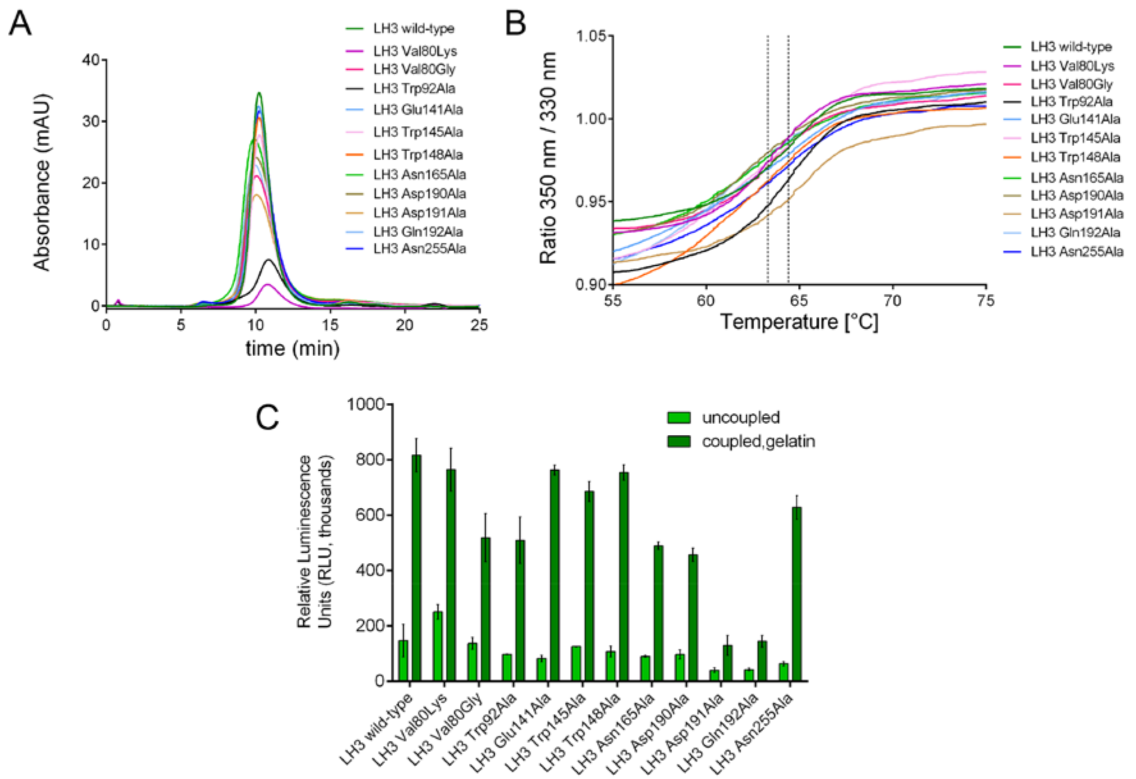


**Supplementary Figure S1: biochemical evaluation of the folding state of LH3 wild-type and mutants.** (A) Analytical size-exclusion chromatography analysis comparing wild-type and mutants LH3. (B) DSF comparing wild-type and mutants LH3. The dashed lines indicate the temperature range incorporating the calculated $T_m$ values for all curves. (C) LH activity comparison for wild-type and mutants LH3. The measurements were performed in absence ("uncoupled") and in presence ("coupled") of gelatin acceptor substrates, as described in (Scietti et al, 2018). Error bars represent standard deviations from average of triplicate independent experiments.

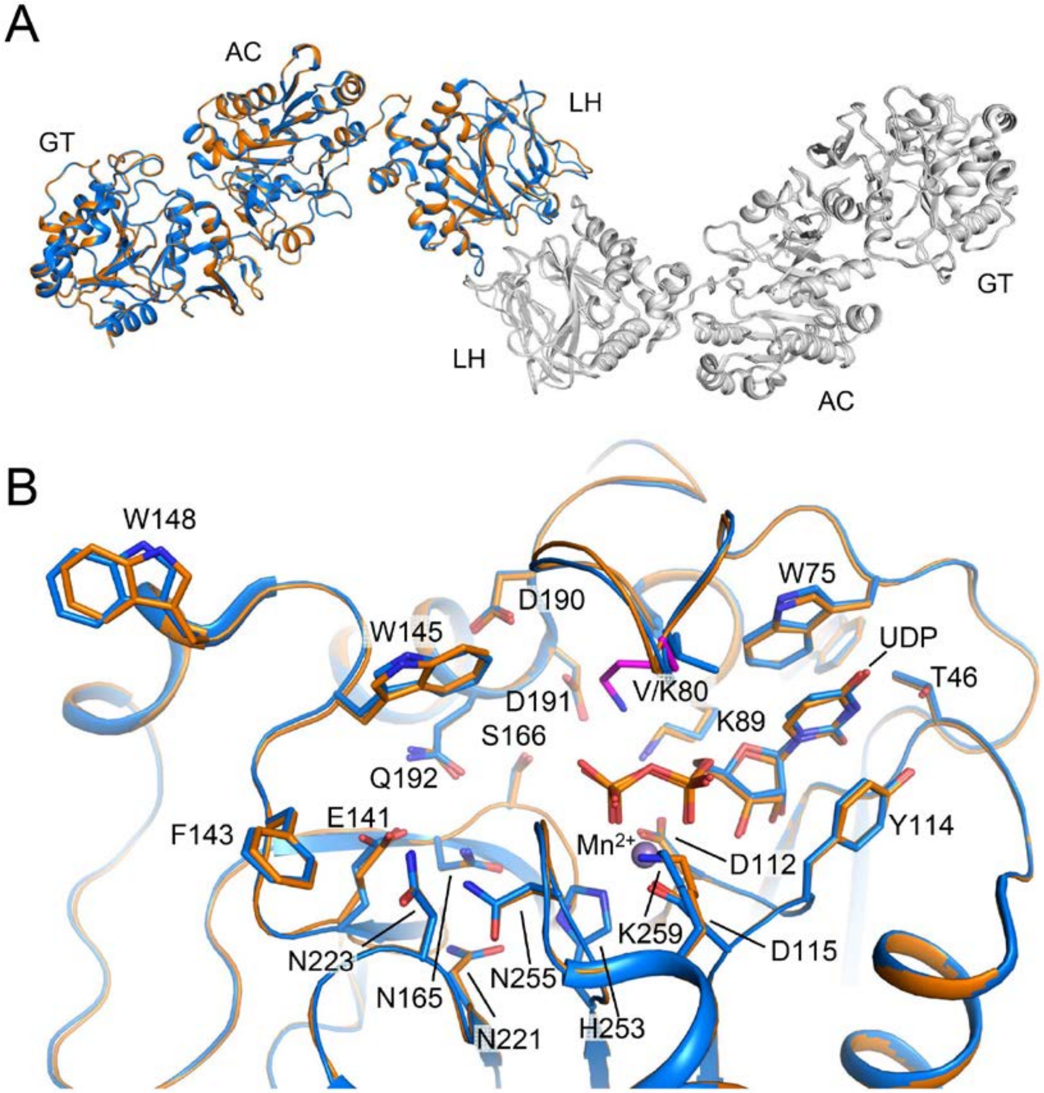


**Supplementary Figure S2: comparison of the LH3 wild-type and the LH3 V80K mutant structures in complex with UDP-glucose.** (A) The overall structure superposition of the wild-type LH3 (blue) and Val80Lys mutant (orange) shows that the two structures are almost identical. (B) Zoomed view of the GT active site of the two structures shown in (A), indicating that the sole difference is constituted by the mutated Val80 to Lys (shown in pink).

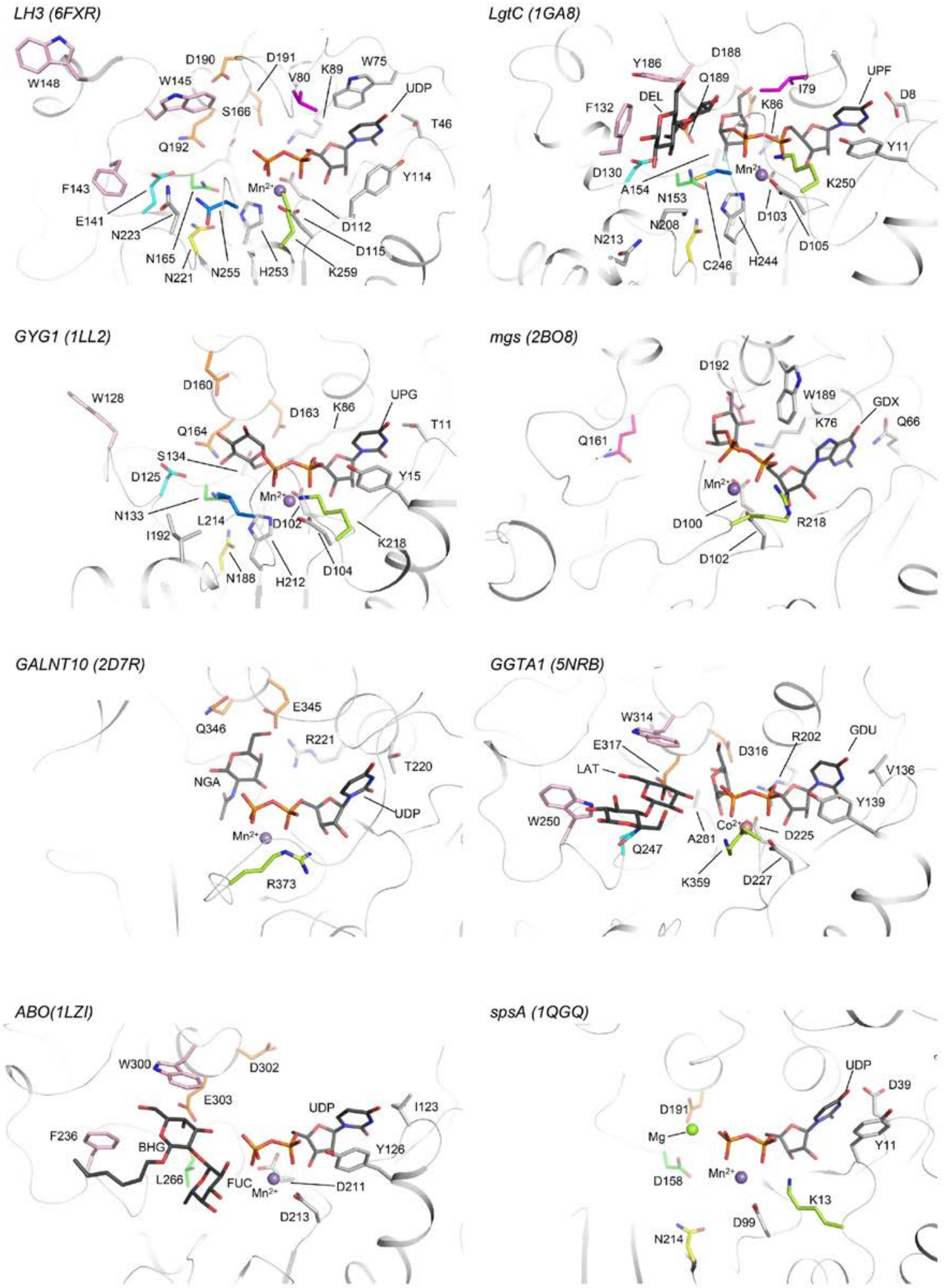
LH3 (6FXR)
W148
D190
D191
V80
K89
W75
UDP
W145
S166
T46
Q192
F143
Mn2+
Y114
E141
N223
D112
N165
D115
N221
N255
H253
K259
LgtC (1GA8)
Y186
D188
I79
UPF
Q189
D8
F132
DEL
K86
Y11
D130
Mn2+
A154
K250
N153
N208
D103
N213
D105
C246
H244
GYG1 (1LL2)
W128
D160
UPG
D163
K86
T11
Q164
S134
Y15
D125
Mn2+
N133
L214
D102
I192
K218
N188
H212
D104
mgs (2BO8)
D192
W189
GDX
K76
Q66
Q161
Mn2+
D100
R218
D102
GALNT10 (2D7R)
E345
Q346
R221
T220
NGA
UDP
Mn2+
R373
GGTA1 (5NRB)
W314
R202
GDU
E317
D316
V136
LAT
Y139
W250
Co2+
A281
D225
Q247
K359
D227
ABO(1LZI)
W300
D302
E303
UDP
I123
F236
BHG
Y126
L266
FUC
Mn2+
D211
D213
spsA (1QGQ)
UDP
D39
D191
Mg
Y11
Mn2+
D158
K13
D99
N214

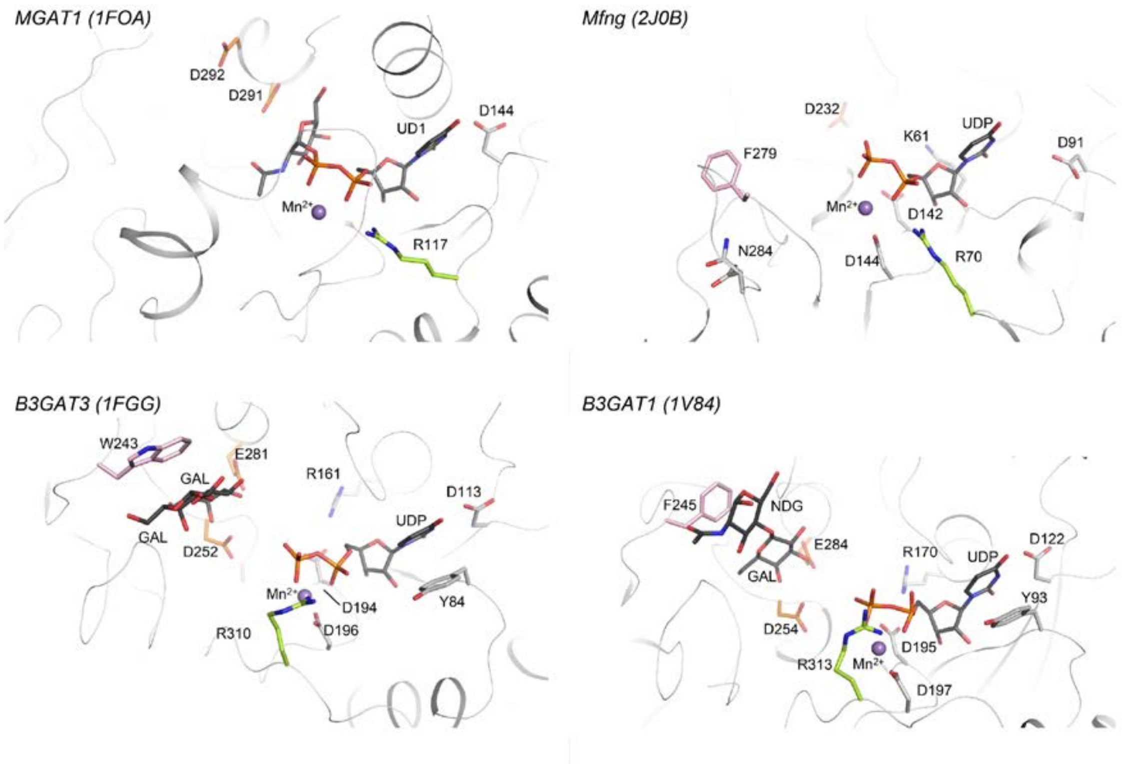


**Supplementary Figure S3: structural comparison between LH3 and other glycosyltranferases.** The conserved aminoacids are shown as sticks; where present, $Mn^{2+}$ cofactor and UDP-sugar are shown as purple sphere and black sticks respectively. Protein name and related PDB ID are indicated for each panel. Colour coding is as in Figure 1A and is maintained throughout the structures allowing to compare the LH3 critical residues with the other structures. For additional information on the proteins indicated in this panel refer to Table 1.

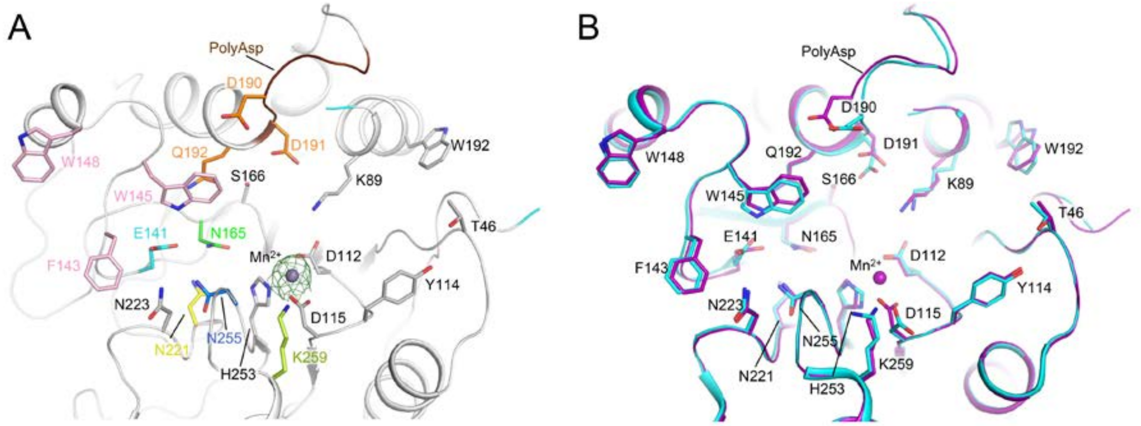


**Supplementary Figure S4: comparison of the GT domains of wild-type LH3 in ligand-bound and ligand-free states.** (A) The structure of wild-type LH3 crystallized in presence of $Mn^{2+}$ and UDP shows clear electron density for $Mn^{2+}$ ($2F_o$-$F_c$ omit electron density map shown as green mesh, contour level 2 σ), but unexpectedly no density is present for UDP. Residue highlight and colouring as in Figure 1A. (B) The comparative superposition of wild-type LH3 GT domain co-crystallized in presence of $Mn^{2+}$ and UDP (purple), and without cofactors (cyan – PDB ID: 6FXK) shows no differences in the conformations of the side chains for the residues delimiting the GalT/GlcT catalytic site.

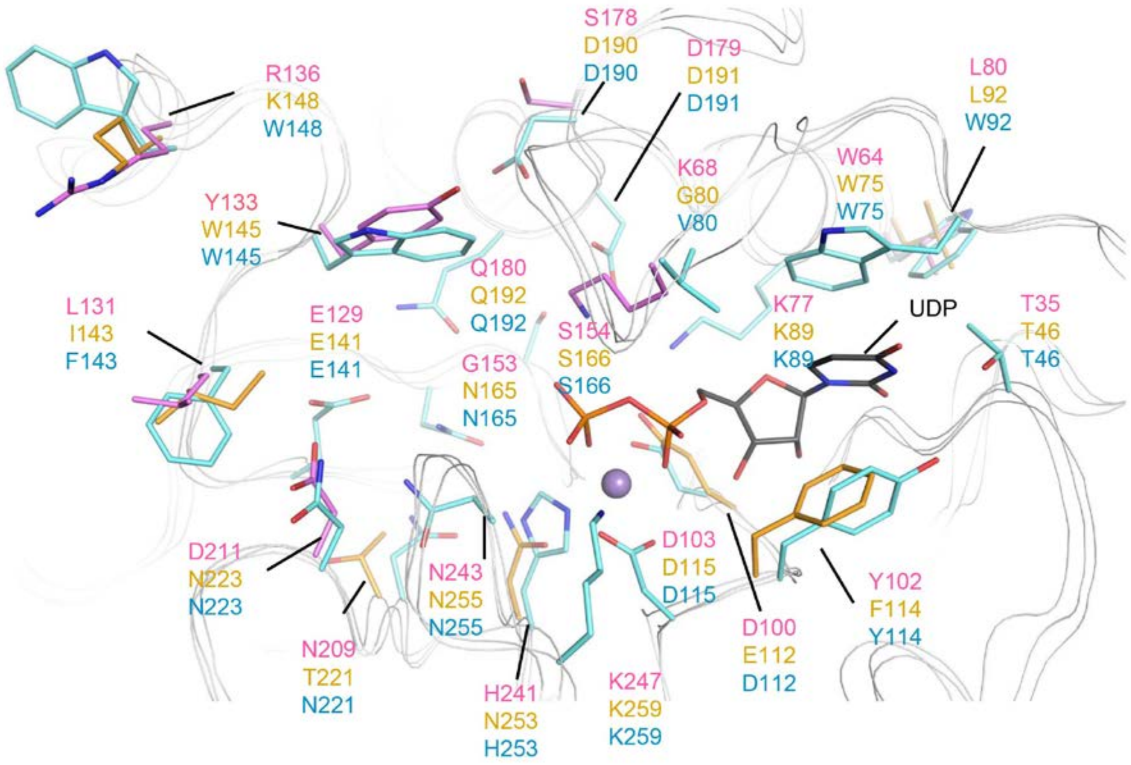


**Supplementary Figure S5: superposition of the LH3 structure and the LH1 and LH2 homology models.** Superposition of the GT site of LH3 in complex with UDP-glucose (PDB ID: 6FXT) and LH1 and LH2 homology models (Scietti et al, 2019) (available at http://fornerislab.unipv.it/SiMPLOD/). Aminoacids identified as part of the LH3 catalytic site are shown as light blue sticks, while the LH1 and LH2 residues which differs from LH3 are shown in pink and orange respectively. UDP moiety (black lines) and $Mn^{2+}$ cofactor (purple sphere) are also shown from the LH3 structure.
